# The conformational ensemble of an intrinsically disordered protein explains peak shapes under DNP conditions

**DOI:** 10.1101/2022.10.22.513343

**Authors:** Jaka Kragelj, Rania Dumarieh, Yiling Xiao, Kendra K. Frederick

## Abstract

Elucidating the conformational preferences of regions of intrinsic disorder in biologically relevant contexts represents a frontier of structural biology. The sensitivity enhancements conferred by DNP enable structural studies of proteins in native contexts by MAS NMR. However, DNP requires low temperatures which results in broad peaks, particularly for for regions of intrinsic disorder. We describe an approach to predict and interpret peak shapes for frozen regions of intrinsic disorder in terms of dihedral angle populations. We demonstrate the method using the protein a-synuclein. This approach can be used to obtain experimental structural restraints for regions of intrinsic disorder in both simplified and biological settings, providing information that eludes characterization by diffraction-based methods as well as solution-state NMR spectroscopy and molecular dynamics due to molecular size limitations.

## INTRODUCTION

Regions of intrinsic disorder are of great importance in biology, yet they evade characterization by many biophysical techniques.^1,2^ Chemical shifts from solution-state NMR are one of the most commonly used parameters in the validation of computationally generated structural ensembles that model regions of intrinsic disorder at atomic resolution.^3^ Because of their accessibility, they are also the primary means of characterizing proteins in in-cell NMR experiments.^4^ However, when proteins are characterized inside cells using solution-state NMR spectroscopy, the interactions with the cellular environment can cause peaks to broaden beyond detection. On the other hand, molecular dynamics (MD) simulations could be useful for modeling intrinsically disordered regions (IDRs) in cellular environments, especially since IDR modeling is becoming more accurate,^5,6^ but the approaches to model a crowded cellular environment are still under development.^7,8^ Yet, biological environments can and do restructure regions that are intrinsically disordered in purified settings.^9^

Dynamic nuclear polarization (DNP) magic angle spinning (MAS) NMR is uniquely poised to report on proteins in their native biological environments at their endogenous concentrations. The signal enhancement with DNP makes it possible to acquire MAS NMR spectra of proteins at low micromolar concentrations in reasonable acquisition times.^9,10^ However, the effectiveness of DNP MAS NMR experiments depends on the experimental conditions and sample preparation.^11–16^ One of the conditions currently required for effective DNP is cryo-temperatures, typically around 100 K.^17^ Low temperatures enable efficient excitation of the nuclei, which are polarized by unpaired electrons introduced into the sample in the form of stable radicals at millimolar concentrations.^15,17^ The increase in experimental sensitivity can be sufficient to enable detection of molecules present at low micromolar concentrations, even in complex biological mixtures.^9,18^

However, low temperatures often have an unfavorable effect on NMR line shapes due to inhomogeneous line broadening from the freezing out of molecular motion.^19–22^ At room temperature, regions of intrinsic disorder sample many conformations on the fast exchange time scale which results in a single peak in the NMR spectrum at the population weighted average chemical shift value.^23,24^ Indeed, deviations from random coil chemical shift value of the peaks are used to uncover conformational biases in the underlying conformational ensembles of intrinsically disordered regions (IDRs).^25,26^ However, at cryogenic temperatures, the dynamic averaging of the chemical shift does not occur, and the static disorder of frozen IDRs results in peaks that are exceptionally broad but not featureless.. It is useful to think of these peaks composed of sub-peaks belonging to multiple conformers that are at the limit of spectral resolution. Because chemical shifts report on local structurethe peak shapes could be used to reconstruct the ensemble of underlying conformations. Thus, analysis of peak shapes obtained under DNP conditions can provide experimental information about the conformational ensemble.

While it has been long recognized that the cross-peak line shapes of frozen samples could serve as quantitative constraints on backbone conformational distributions in terms of α- helical and extended conformations,^27–29^ recent technical advances for DNP MAS NMR such as higher operating magnetic fields and cryogenic probes that support faster spinning rates potentially reduce the sources of homogenous line broadening enough to potentially enable useful deconvolution of the experimental spectra. Indeed, for highly ordered systems measured under conditions that differ only in experimental temperature, the line widths at ambient and cryogenic temperatures are similar.^19,30^ Thus, the broader line widths for biological samples at cryogenic temperatures come from the freezing out of the small excursions around a local minimum that are not detected in room temperature experiments. This effect is exaggerated for loops and termini because the peak reports on all the sampled conformations with peak intensities related to the relative populations of each conformation in the ensemble. In such cases, knowledge of the expected peak shapes for a completely unrestrained amino acid in a polypeptide chain would be useful to interpret the degree of structural restraint at a particular site. Any deviations of the experimental spectra from the expected peak shapes for a completely disordered region indicate conformations that are preferred or disfavored. Better approaches for analyzing peak shapes in terms of structure combined with new technologies^31–33^ and biochemical approaches to sample preparation, such as the segmental or specific isotopic labeling of proteins^34–35^ opens up the possibility of examining one, or a few sites, in the context of a full-length protein.

In this work, we examine the spectra of frozen samples of the protein α-synuclein as an intrinsically disordered monomer as well as in an α-helical membrane bound form^[36–39^ and a β- sheet rich amyloid form^40-44^ under conditions with reduced homogenous line broadening that are representative of modern commercial DNP spectrometers. We build structural ensembles of intrinsically disordered regions using a statistical coil library, determine the chemical shifts for each member of the ensemble using chemical shift prediction software and simulate the spectra for each of the amino acids when it is intrinsically disordered. We annotate these spectra with the corresponding φ/ψ-region to indicate which conformations give rise to the different components of these compound peaks.

## RESULTS AND DISCUSSION

### Different conformations of α-synuclein are distinguishable under DNP MAS NMR conditions

To determine how well we can experimentally distinguish between β-sheet, random coil, and α-helical conformations, we collected DNP spectra of α-synuclein in three different conformations; a β-sheet rich amyloid fibril form,^40^ a frozen sample of disordered monomeric protein, and an α-helical-rich nanodisc-bound form.^29,36^ To eliminate chemical shift ambiguity, α-synuclein was isotopically labeled at only the threonine residues (Figure 1A). As expected, the ^13^C-^13^C DARR spectra of specifically threonine-labeled α-synuclein had cross peaks for the threonine C’-C^α^ & C’-C^β^ correlations (Figure 1B) and C”-C^γ^ & C^β^-C^γ^ correlations (Figure 1E). To facilitate the examination, we calculated projections for each peak and we annotated the projections of spectra with the tabulated average chemical shift values for threonine C’, C^α^, and C^β^ sites in theα-helical, random coil and β-sheet conformations.^45^ The tabulated average chemical shift values for each secondary structure (Figure 1; crosses and filled circles) indicated well where we can expect the peak centers.

**Figure 1:**
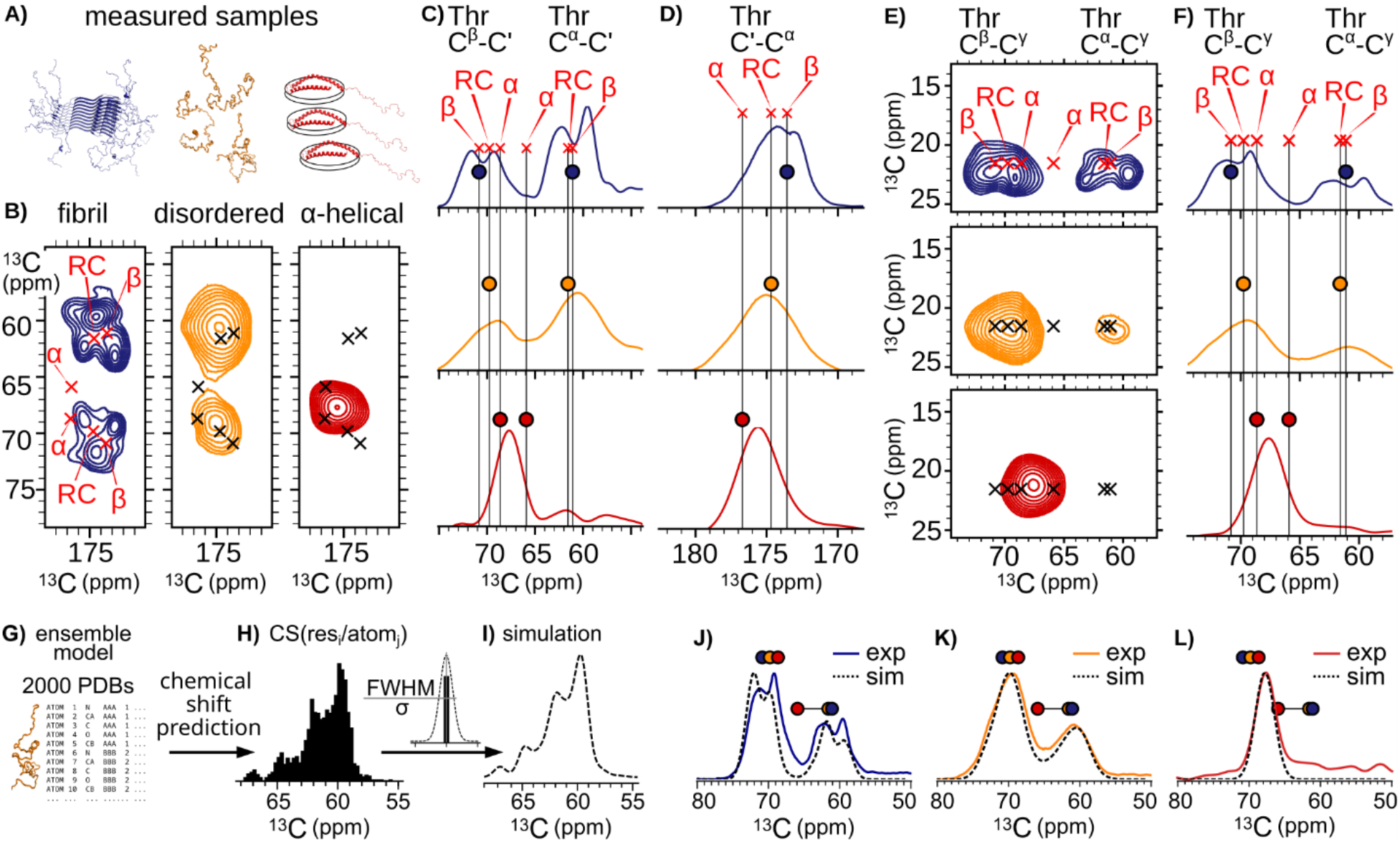
Chemical shift changes between different α-synuclein conformations and simulation of peak shapes. A) models representing the three samples: fibrillar α-synuclein (blue; PDB: 2N0A), α-synuclein in solution (orange), α-synuclein bound to nanodiscs (red). B) Aliphatic-carbonyl region of ^13^C-^13^C DARR with chemical shifts annotations for β-strand, random coil, and α-helix from Wang & Jardetzky, 2002. C) Projections of B) on C^α^/C^β^-axis. D) Projections from B) on C’-axis. E) Region of ^13^C-^13^C DARR containing correlations for C^γ^-C”and C^γ^-C^β^ of threonines. F) Projections of E) on C^α^/C^β^-axis. G, H, I) Schematics showing how we simulated the statistical coil ensemble for the disordered α-synuclein and how the peak shapes are simulated from the predicted chemical shifts. J, K, L) Comparison of simulated chemical shifts with the projections of the ^13^C-^13^C DARR spectra for fibrillar α-synuclein (blue), α-synuclein in solution (orange), α- synuclein bound to nanodiscs (red).

In the case of the fibrils and the disordered state, the C^α^ and C^β^ regions were easily distinguishable while in the case of the α-helical form of α-synuclein, these regions had only one apparent peak due to similarity of the chemical shift values of the threonine C^α^ and C^β^ for α-helical conformations and the homogenous line widths under the experimental conditions (Figure 1; red color). This overlap, resulting in a merged peak, is problematic for the interpretation of secondary chemical shifts in α-helical proteins such as transmembrane proteins measured under DNP conditions.

Interestingly, the peaks of the disordered α-synuclein had each only one maximum, which seemingly conflicted with the expectation of several peaks of different frozen conformations. However, intrinsically disordered regions have a high population of the P_II_ and δ conformations with distinct chemical shifts that fall between those for α-helical and β-sheet structures (i.e. around the random coil value).^24,46^ A combination of P??, δ, α-helical conformations, and β-strand conformations with enough heterogenous and/or homogenous peak broadening could result in a broad peak with the peak maximum close to the tabulated random coil value. We thought that it would be interesting to inspect the origin of the random coil peak shape more closely.

### Obtaining peak shapes from chemical shift predictions from a statistical coil ensemble

To study the peak shapes in more detail, we simulated the expected peak shapes for the 10 threonines in the disordered, nanodisc-bound, and fibril forms of α-synuclein using structure-based chemical shift prediction. To simulate the disordered α-synuclein, we created an ensemble of 2000 PDB structures (Figure 1G). We created the structures using flexible meccano which builds disordered polypeptide chains based of a given sequence. During the building process, the φ/ψ dihedral angles of each residue are selected from a restricted coil library.^[24,47–49]^ The restricted coil library contains amino statistics of φ/ψ samping preferences specific to each amino acid and describes the sampling of IDRs accurately enough for comparisons with experimental data.^[23,46,47]^ We therefore refer to this ensemble representing disordered α-synuclein as a statistical coil ensemble (i.e. sampling of conformations follows the statistics in the restricted coil library). We then predicted the chemical shifts for each conformer in the ensemble using a static chemical shift predictor PPM_One (Figure 1H).^50^After obtaining the distributions of the predicted chemical shifts, we added 1.9 ppm of line broadening to account for the homogenous line broadening under these experimental conditions (Figure 1I). The fibrillar form and the nanodisc-bound form have both structured and disordered regions, and our predictions took both into account. For the fibrillar form of α- synuclein, we used an experimentally-determined structure with the disordered N- and C- tails modeled as statistical coil.^40^ The nanodisc-bound form was modeled as a continuous α-helix spanning residues 1 – 93 and the disordered C-tail of the nanodisc-bound form was modeled as a statistical coil. The simulated spectra captured the peak shapes and centers of experimental spectra better than the tabulated chemical shift values from work of Wang & Jardetzky^45^(Figure 1K,L, Figure S1). The centers of the simulated peaks for the fibril form differed slightly from the experimental spectra (Figure 1J, Figure S1). This discrepancy could arise from dynamics and flexibility not represented in the solved structure; of the ten threonines in α- synuclein, in this fibril form, eight are in the rigid amyloid core (five in the middle of beta sheets and three at the end of sheets or in turns) and two are in a region that was not visible in the room temperature solid state NMR spectra.^40^ Indeed, these small disagreements strongly suggest that the mobile regions of the fibril form at room temperature sample a biased subset of conformations. In summary, the improved agreement of the simulated and experimental spectra demonstrates that our simulation framework captured the major features of the NMR spectra, particularly for a frozen intrinsically disordered form of α-synuclein.

The use of a chemical shift predictor that does not employ a heuristic to intuit dynamics and alter the predicted chemical shift, such as PPM_One,^50^ was critical for obtaining good agreement with the projection of the disordered α-synuclein. Other chemical shift predictors, such as SPARTA+,^51^ average the chemical shifts for loops and disordered regions toward the random coil values. This results in a narrower chemical shift distribution for the disordered α- synuclein, and consequently, the simulated spectrum does not capture the full breadth of sampled chemical shifts (Figure S2).

### Mapping contributions of different conformations to the final peak shape

Having established that the simulations can recapitulate the shape of the projections, we investigated how the individual φ/ψ populations contribute to the final peak shapes in the experimental spectra. Starting from the ensemble model of disordered α-synuclein, we binned the conformers of each threonine according to the φ/ψ angles with a bin size of 5° (Figure 2A). We represented the average C^α^ chemical shift value within each bin with a color, while the bar heights represent the population in each bin (Figure 2B,C). We classified the regions of the Ramachandran plot (Figure 2D) following the naming convention defined by S.A. Hollingsworth and P.A. Karplus.^52^ We defined regions of the Ramachandran plot such that all bins are assigned to a φ/ψ-region. The region normally associated with the chemical shift of α-helix (Figure 2D,E) was defined to contain conformations that could form a repeating α-helix structure.^53^ The region with positive ψ-angles, usually associated with extended conformations, was divided into polyproline II region (P_II_, sky blue), ζ- and γ’-region (light cyan), and the region representing conformations found in β-strands (dark blue). Here we refer to region with positive φ angles as δ’ or α?_. This conformation is most often occupied by glycines or asparagines when they are part of turns. The δ-region (orange) occupies the space around the α region. We mapped the φ/ψ-regions of disordered α-synuclein to the chemical shift distributions (Figure 2F,G,H). To do this, we added the φ/ψ information to each predicted chemical shift. We visualized the data by color-coding the chemical shift distributions according to the φ/ψ-region of origin which highlighted how different regions contribute to the overall chemical shift distribution and peak shape.

**Figure 2:**
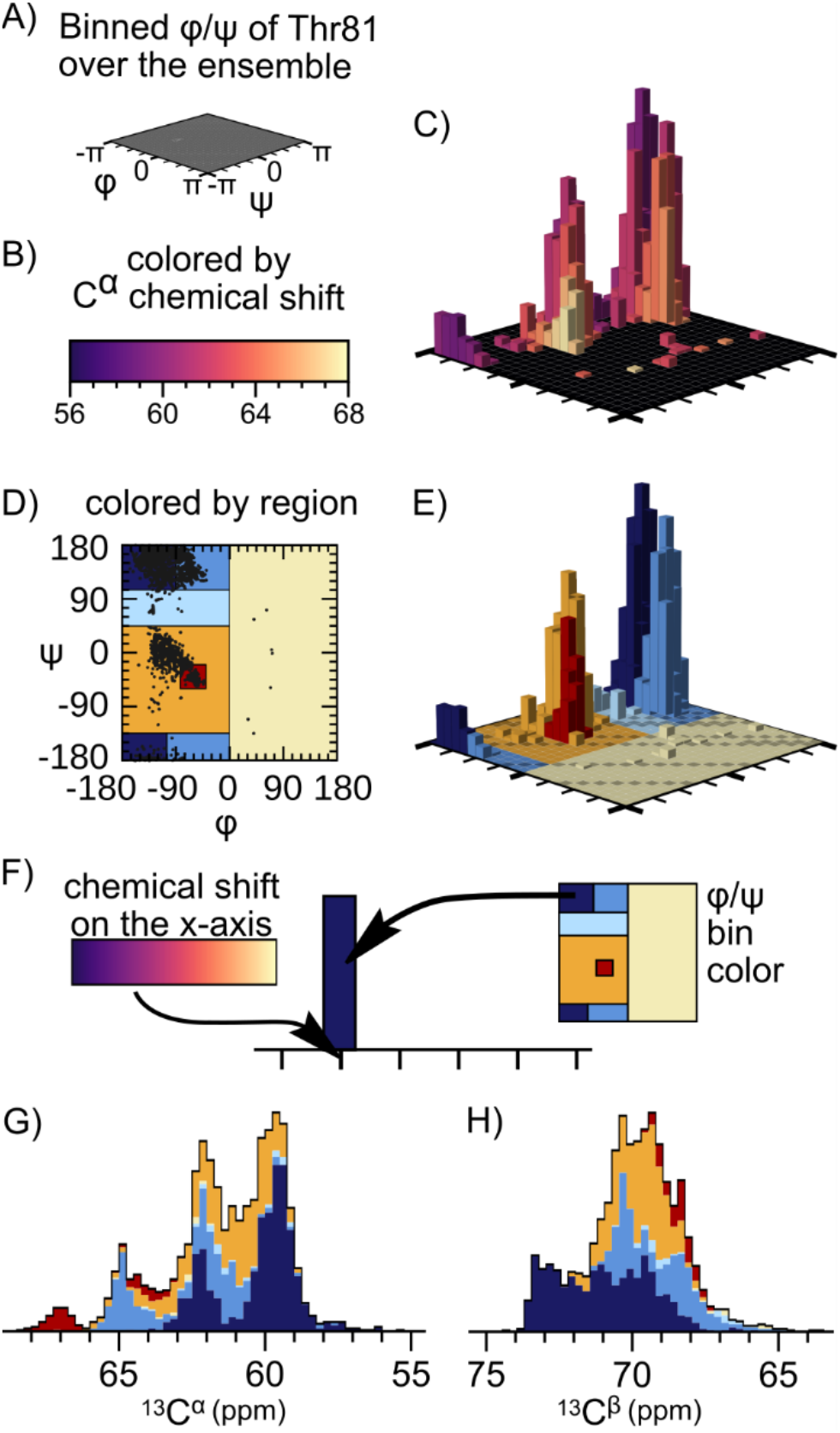
Mapping of φ/ψ-regions sampled in the statistical coil ensemble to simulated chemical shift distributions. A) Legend for the Ramachandran plots C and E. B) Color legend for bar plot C. C) Conformers of threonine 81 from the statistical coil ensemble of disordered α- synuclein were binned into 5° bins according to their φ/ψ angles. The colors of the bars indicate the average chemical shift values in each bin. The bar height represents the conformer count for that bin. D) Definitions of regions based on φ/ψ angles. E) Conformers of threonine 81 from the statistical coil ensemble of disordered α-synuclein binned into 5° bins according to their φ/ψ angles and colored according to φ/ψ-region definitions from D. F) A scheme displaying how each bar in G and H simultaneously represents the average chemical shift of the bin and the φ/ψ- region of origin. G,H) Chemical shift predictions colored according to the φ/ψ-region of origin.

Examination of chemical shift distributions for threonine revealed that non-α and non-β conformation were significantly populated. There were four subpeaks in the C^α^ chemical shift distribution (Figure 2G). The small peak downfield was the result of the conformers in α-helical conformation. The largest peak up-field was mainly the result of the conformers in β-strand conformation. The presence of the two middle non-α- and non-β-peaks explains why the experimental spectrum of disordered α-synuclein differs from those of the α-helical and fibrillar samples. While the two middle peaks in the experimental spectrum were not resolved due to line broadening, their presence explains why the experimental spectrum of α-synuclein had a single broad peak centered at the random coil value.

### Limitations of the statistical coil model

The statistical coil library quantitatively described the conformations in the ensemble and the experimental spectrum. However, it is well appreciated that the populations of φ/ψ- regions in statistical coil libraries might deviate from the distributions obtained by detailed experimental studies.^[23,24,46,54]^ Additionally, the sampling of our ensembles was almost neighbor independent. Rudimentary steric restraints were taken into account during model building but the conformational preferences that rely upon context and long-range interactions are not well-captured using the statistical sampling method. This feature of the statistical sampling approach to generate ensembles was evident when we examined the simulated the peak shapes for individual threonine residues in our statistical coil ensemble. The simulated peak shapes for all the threonines in α-synuclein were very similar (Figure 3). The only minor deviation was in the simulated peak shapes for the carbonyl carbon which was centered around 174 ppm for all sites except those that are followed by a glycine in the primary sequence which were centered around 175 ppm. Therefore, the predicted peak shapes using this statistical approach are for an idealized region of intrinsic disorder devoid of any secondary structure propensities. Spectra predicted from such model can be used to identify sub-peaks belonging to each φ/ψ-region. This stems from the fact that the deviations of φ/ψ-region populations affect the intensities of sub-peaks while sub-peak positions are more likely dependent on the accuracy of the chemical shift predictor.

**Figure 3:**
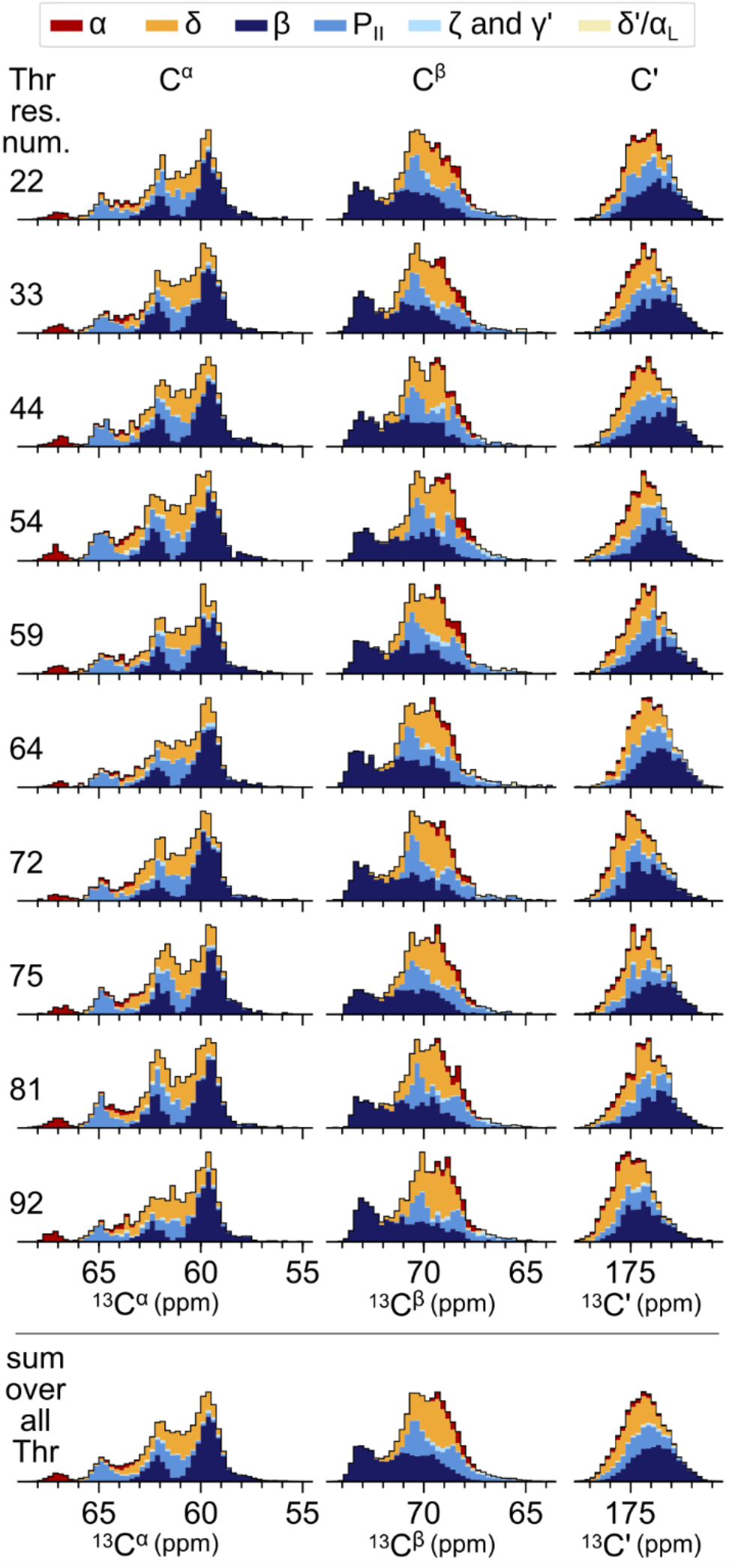
Prediction of chemical shifts for every threonine of the α-synuclein statistical coil ensemble. Chemical shift predictions for threonines of the disordered α-synuclein ensemble. Color indicates how populations of φ/ψ-regions translates into peak shape. The bottom most histogram represents the sum over all threonines in the α-synuclein sequence.

### Assigning peaks in ^13^C-^13^C correlation experiments

To determine whether our simulated chemical shift distributions can be used to assign cross-peaks in 2D correlation experiments, we compared the simulated peak shapes for a frozen region of intrinsic disorder to the experimental peak shapes. To do so, we examined the peak shapes of the threonine C”-C^v^ and C^β^-C^v^ correlations in a frozen sample of α-synuclein. Because the threonine C^v^ chemical shift is not very sensitive to changes in secondary structure,^55^ identification of the φ/ψ-regions of threonine C^α^-C^v^ and C^β^-C^v^ cross-peaks relied on the C^α^ and C^β^ chemical shift information independently, and the sub-populations corresponding to the different Ramachandran regions were not well resolved (Figure S3). We then examined the valine C^α^-C^β^ cross peak because this cross-peak has been studied before in the same context.^29^ We prepared a sample of disordered α-synuclein labeled with [2-^13^C]-glucose following previously published protocols.^29^ This labeling scheme results in a single valine C^α^-C^β^ cross peak in the aliphatic region of ^13^C-^13^C correlation spectra. The peak for the C^α^-C^β^ correlation had a curved shape because both the C^α^ and C^β^ chemical shifts are sensitive to secondary structure (Figure 4A).

**Figure 4:**
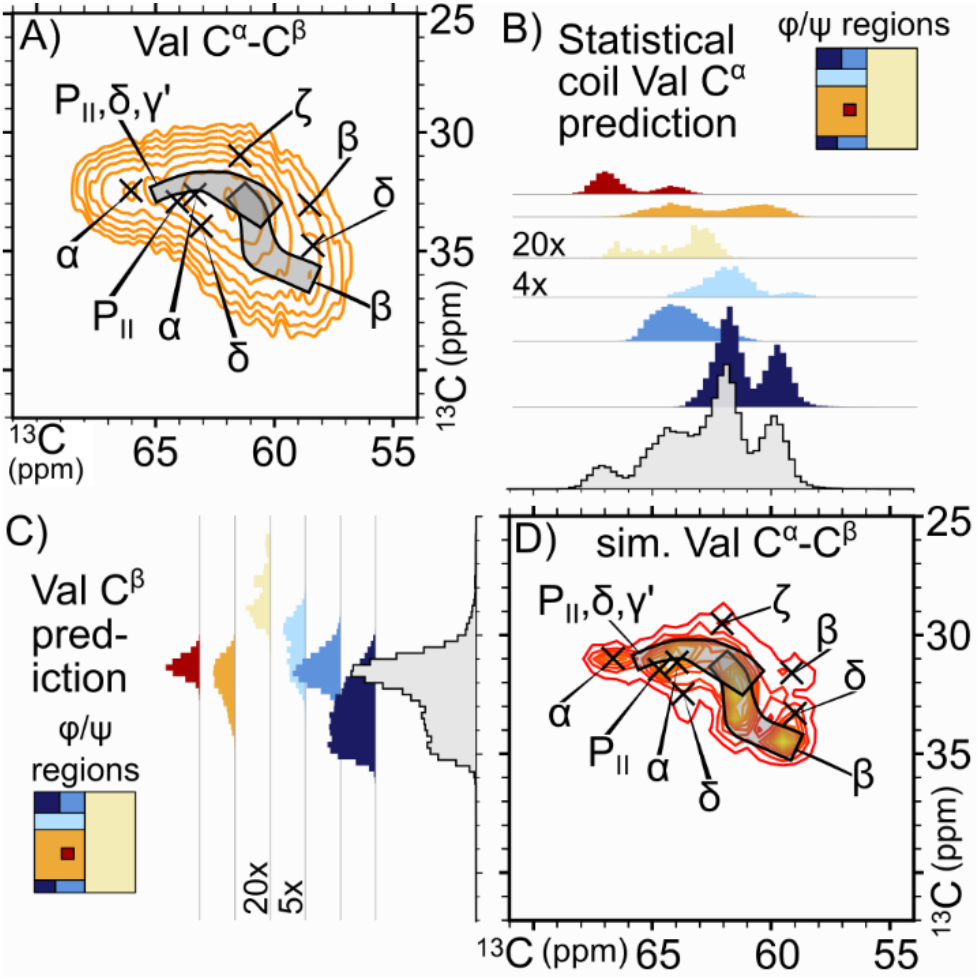
Conformational annotations from the simulated spectra mapped onto experimental spectra for the valine C^α^-C^β^ cross-peak of frozen disordered α-synuclein. A) The valine C^α^-C^β^cross-peak of a sample of frozen disordered α-synuclein was annotated with the conformations corresponding to φ/ψ-regions that were determined from the simulated spectra. Rectangles are used to indicate φ/ψ-regions when the spread of the chemical shift values is too large to be represented by a single cross. B and C) Predicted chemical shift distribution for the (B) valine C^α^atom and (C) valine C^β^ atom plotted as sub-peaks specific to the φ/ψ-regions for α (red), δ (orange), β (dark blue), P_II_(blue), ζ & y’ (light blue), and δ/α_L_(yellow) conformations as well as the sum (gray). D) Contour plot of the simulated valine C^α^-C^β^ cross-peak of the Ala_25_-Val-Ala_25_ statistical coil ensemble. The φ/ψ-regions of the 2D cross-peaks were annotated with the aid of the 1D sub-peak histograms from B) and C).

Next, we wanted to generalize our chemical shift predictions so that the results would be applicable to other proteins with intrinsically disordered regions. We constructed an ensemble for the host-guest peptide Ala_25_-Val-Ala_25_. To validate this approach, we compared the results of the host-guest peptide simulations with the experimental C^α^- C^β^ correlations of valines in α-synuclein. We predicted the chemical shifts of valine as described for threonine above, except that this time we also plotted a 2D histogram of the chemical shift distribution (Figure 4D). The 1D distributions of the C^α^and C^β^ chemical shift distributions (Figure 4B,C) served as a guide for assigning the φ/ψ-regions of the experimental valine C^α^-C^β^ cross peak (Figure 4A).

In the assigned valine C^α^-C^β^ cross peak, the α population was well separated from the other populations. The β population occupies the opposite side of peak although with more spread, which is potentially a result of the non-natural rectangular definitions of φ/ψ-regions used in this treatment.^52^ The correlations for the P??population also partially overlap with other populations, although it is interesting to note that this population occupies a well-identifiable narrow region of the compound peak. The δ population, on the other hand, maps to a broader region of the compound peak (Figure 4B,C,D). The γ’, ζ, populations are very small and, for valines, the δ’/αL is negligible.

### Prediction of correlations for each amino acid

The approach using host-guest peptides can be used to predict peak shapes for other amino acids as well. We constructed a collection of Ala25-Guest-Ala25 peptides, where Guest is one of the 20 amino acids (Figure 5). The peak shape predictions can then be compared to the spectra of any protein. We did not make predictions for cysteines because the statistical library does not contain information on their sampling. We performed predictions for C^α^-C^β^correlations (Figure 5) as well as for C^α^-C’, C^β^-C’, N-C^α^/C^β^ (Figure S4 – S11). Assignment of Ramachandran sub-populations was performed with the aid of the 1D chemical shift distributions. We found some similarities in the positions of φ/ψ-regions relative to one another when comparing peak assignments for different amino acids. The chemical shift of the α population was separated from the rest of the conformations for most amino acids and the chemical shifts of the β population were on the opposite side of the predicted compound peak. In contrast to the chemical shifts of the α-helical conformations, which typically populated a single well-defined, intense peak, the chemical shifts from the β-region occupied a wider part of the compound peak, spreading from the periphery to the center. The chemical shifts of the ζ, γ’, and δ’/αL populations were often located towards the periphery of the compound peak (e.g., at the top of the peak for the peaks above diagonal in ^13^C-^13^C correlation spectra). The chemical shifts of the P??, δ and β-regions were the most difficult to separate because the chemical shifts that correspond to these areas of Ramachandran space are often somewhat degenerate. However, each of these regions had some distinguishing features. The P??region typically formed one (or more) well-defined, intense peak(s). The δ population typically formed a curved or diagonal shape which often overlapped with P??peaks. The β-region also typically formed a curved or diagonal shape which was distinct at the periphery of the peak but overlapped with either (or both) of the P??and δ regions near the center of the compound peak. Although the φ/ψ-regions sometimes overlapped, different areas of the compound peak can be assigned to one (or a few) different conformations.

**Figure 5:**
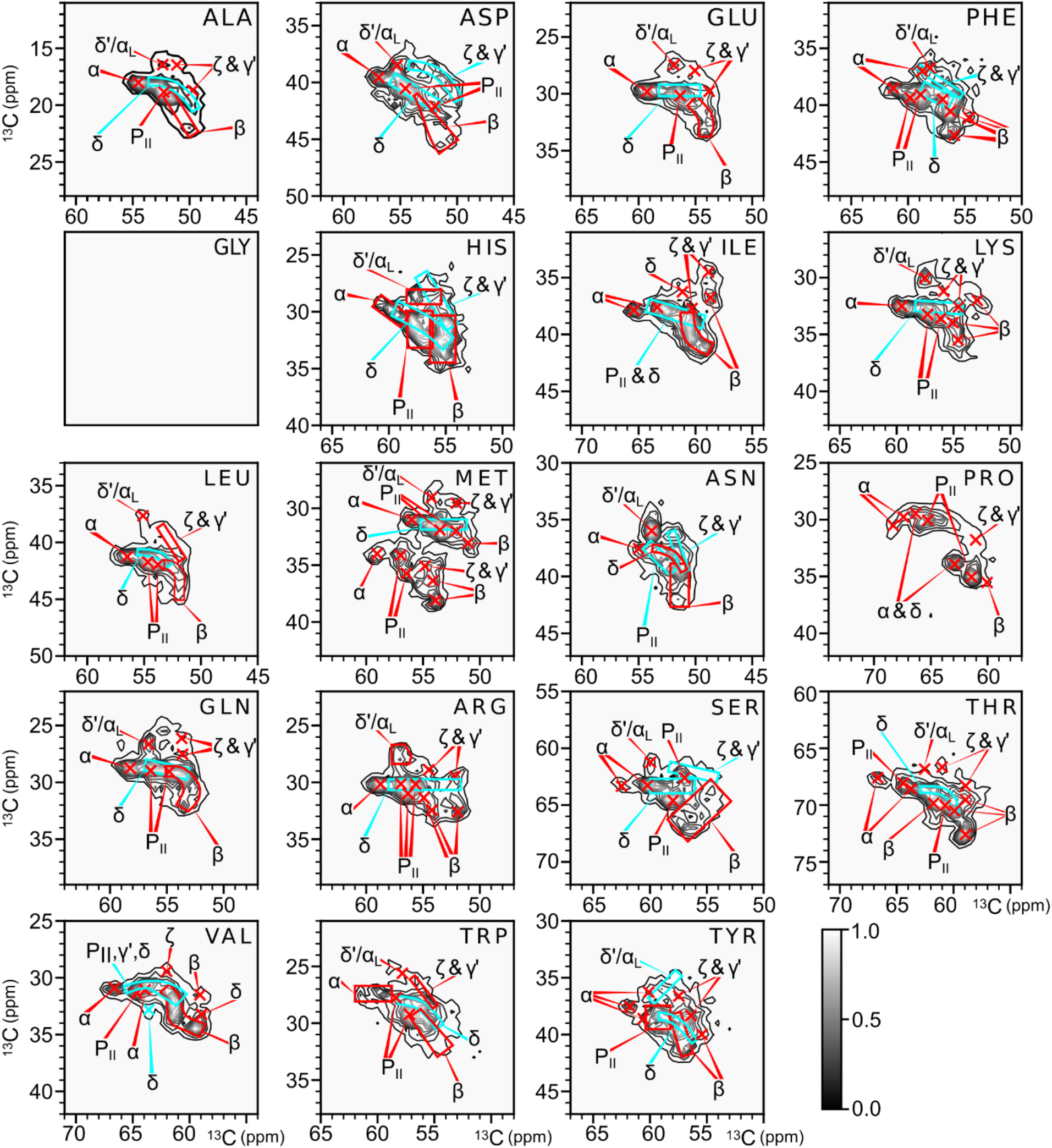
Simulated peak shapes for the C^α^-C^β^ cross peaks for the guest amino acids in the Ala_25_-Guest-Ala_25_ statistical coil ensembles. Plots are ordered according to the single letter amino acid code. The φ/ψ-regions were assigned with the help of 1D chemical shift distributions.

The position of φ/ψ-regions relative to one another is conserved for most amino acids. However, the conserved relative distribution of assigned regions does not mean that the peak shapes must be identical, since the shape of compound peaks is determined by the relative population of the different conformers. For example, aspartate and valine had an almost identical arrangement of assigned regions but aspartate has a high α and low β population, whereas in the case of valine, the peaks of the β population were the most intense. This resulted in a simulated compound peak of aspartate having a diagonal character because the α, δ, P??, and δ’/αL regions were highly populated. In contrast, the simulated compound peak for valine was curved because all the regions for valine were more equally populated than they were for aspartate. Likewise, the simulated compound peak of asparagine, which has a high δ’/αL population, differs from that of both aspartate and valine despite the similarity of the relative distribution of φ/ψ-regions to those of aspartate and valine. The peak shape is defined not only by the distribution of φ/ψ-regions relative to one another but also by the relative population of those regions. Thus, the peak shape of a frozen intrinsically disordered regions reports quantitatively on the relative populations of the sampled conformations and deviations from the predicted peak shape can highlight conformational propensities in the sampled ensemble.

## CONCLUSIONS

The established framework for predicting peak shapes of intrinsically disordered regions under DNP conditions will support future studies of frozen regions of intrinsic disorder. Here, we simulated peak shapes using structural ensembles. We experimentally validated these simulations by comparing the simulated and experimental spectra of threonine and valine in frozen samples of intrinsically disordered α-synuclein. We extended this approach to simulate peak shapes for all the amino acids. Because φ/ψ angles and chemical shifts are correlated, we annotated the different regions of the simulated peaks by the corresponding region of Ramachandran space. The assigned regions (Figure 5) and peak shape predictions (Figure S4 - S11) can be therefore used to guide interpretation of ^13^C-^13^C and ^13^C-^15^N correlation spectra of frozen samples.

Here, we simulated peak shaped using structural ensembles that were generated using a statistical coil model. Simulation of peak shapes from more accurate structural ensembles - like those from MD trajectories rather than simple statistical coil ensembles - will capture the near neighbor effects and long-range interactions that alter secondary structural propensities and result in better predictions. Nonetheless, comparison of experimental spectra to simulated spectra of idealized IDP ensembles devoid of secondary structure propensities presented here will highlight sites with transient structurations. Furthermore, it will be possible to build realistic and accurate IDP ensembles by developing an approach for analysis of peak shapes in terms of φ/ψ-regions’subpopulations. This approach could be particularly powerful for the study of regions of intrinsic disorder in environments that are difficult to approximate using molecular dynamics, such as in liquid-liquid phase condensates or for proteins inside cells.

## ASSOCIATED CONTENT

Supporting Information includes extended methods and supplemental figures S1-S11.

## AUTHOR INFORMATION

### Authors

Jaka Kragelj - Department of Biophysics, UT Southwestern Medical Center, Dallas, Texas 75390-8816, United States;

Rania Dumarieh - Department of Biophysics, UT Southwestern Medical Center, Dallas, Texas 75390-8816, United States;

Yiling Xiao - Department of Biophysics, UT Southwestern Medical Center, Dallas, Texas 75390-8816, United States;

## Author Contributions

J.K. and K.K.F conceived the project. J.K. and R.D. prepared the samples. J.K. and Y.X. acquired the spectra. J.K. performed the data analysis and simulations. J.K and K.K.F. wrote the manuscript.

## Notes

The authors declare no competing financial interest.

## ACKNOWLEDGMENTS

This work was supported by grants from the National Institutes of Health (NS-111236), the National Science Foundation (1751174) the Welch Foundation (I-1923_20200401) to K.K.F.

## ABBREVIATIONS

NMR: nuclear magnetic resonance
MAS: magic angle spinning
DNP: dynamic nuclear polarization
IDR: intrinsically disordered region
MD: molecular dynamics.

