## Supplemental Methods for "The conformational ensemble of an intrinsically disordered protein explains peak shapes under DNP conditions"

#### Protein expression and isotopic labeling

The *E. coli* strain BL21 DE3 was used to express isotopically labeled wild-type α-synuclein. The protocol for specific isotopic labeling of threonines was based on previously published protocols (Rime Kerfah 2014, Velyvis et al. 2012). First, the *E. coli* cells were grown in 4 liters of M9 supplemented with natural abundance threonine (50 mg/ml), α-ketobutyrate (100 mg/ml), and natural abundance glycine (500 mg/ml). Cells were grown at 37 °C until the $OD_{600}$ of the culture reached 0.6 – 0.8. The *E. coli* cells were then harvested, washed once with 1x M9 salts (Marley et al. 2001), and transferred to 4 liters of M9 supplemented with $^{13}C,^{15}N$-threonine (50 mg/ml), α-ketobutyrate (100 mg/ml), and natural abundance glycine (500 mg/ml). Cells were then grown at 37 °C until they reached $OD_{600}$ of 1.5. Expression of α-synuclein was induced with addition of 1 mM IPTG. Cells were harvested after 3 hours of expression at 37 °C.

Previously published protocol (Uluca B. et al. 2018) was followed for the expression of α-synuclein labeled at the valine $C^{\alpha}$ and $C^{\beta}$ atoms. Briefly, *E. coli* cells were grown in 2 L natural abundance M9 medium at 37 °C until the OD600 of the culture reached 0.6. Cells were then harvested, washed once with 1x M9 salts (Marley et al 2001), and transferred to 0.5 L M9 minimal media with 4 % (w/v) D-glucose-2-$^{13}C$ (CIL Inc., USA). After a 30 min incubation at 37 °C, the cells were induced with 1 mM IPTG. Cells were harvested after 3 hours of expression at 37 °C.

### Protein purification

Purification was performed following the protocol reported by Barclay, A. M. et al. 2018. Cells that were frozen after the expression were resuspended in lysis buffer containing a detergent (20 mM Tris pH 8.0, 1 mM EDTA, 0.1% Triton-X) and incubated for 30 minutes at 37 °C. After the incubation, DNA and RNA were digested by adding 2.5 µl Omni Nuclease and 2.5 µl DNase while supplementing the solution with 10 mM MgCl2 and 10 mM CaCl2 final concentration. After 1 hour incubation at 37 °C the excess metal was chelated by addition of 5 mM EDTA. Insoluble cell debris was separated from the supernatant by centrifuging at 4,000 xg for 15 minutes. The concentration of NaCl in the supernatant was adjusted by mixing 6 parts of supernatant with 1 part of 5 M NaCl. The supernatant was heated above 90 °C for 10 minutes in a water bath, after which the supernatant is cooled down in a room temperature water bath and on ice. The proteins that precipitated during the boiling step were pelleted in SS-34 rotor tubes, centrifuged at 47,800 xg for 20 minutes at 4 °C. The soluble supernatant containing α-synuclein was precipitated by addition of cold ammonium sulfate at 1:1 ratio (final concentration of 50 % saturated ammonium sulfate at 4 °C). The solution was stirred overnight in the cold room. The following day, the precipitated fraction was separated from the supernatant by centrifuging at 4000 xg for 20 minutes at 4°C. The pellet was dislodged with a spatula so that the clumps could be collected on a 0.22-micron bottle-top filter by pouring the supernatant over the filter. After changing the collection bottle, the pellet trapped on the filter was dissolved in IEX buffer A (20 mM Tris pH 8.0, 20 mM NaCl). The solution was loaded onto a Q-Sepharose column and α-synuclein was eluted with a salt gradient of 0-500 mM NaCl. The a-syn containing fractions were pooled and concentrated to 1 mM and then run over a size exclusion column (Superdex 75 Increase HiScale 26/40, 40 cm). Size exclusion separates α-synuclein from its degradation product with an approximate molecular weight of 11 kDa. Pooling only the purest fractions from size exclusion one can achieve purity of > 98 %.

### Preparation of frozen intrinsically disordered monomers

The stock α-synuclein solution at 1 mM was diluted in D2O to result in 10 mM sodium phosphate buffer at pH 7.0 at a 12:88 ratio of $H_2O$:$D_2O$. Glycerol was added to a final concentration of 15 % d8-12C-glycerol resulting in a 10:75:15 ratio of $H_2O$/$D_2O$/$d_8$-$^{12}C$-glycerol. AMUPol was dissolved in the sample to result in final concentration of 10 mM AMUPol. The sample was transferred to a rotor with a pipette and plugged before being stored at -80 °C until measurements. Prior to measurement samples were briefly warmed to room temperature to mark and cap the rotor. Room temperature rotors were inserted into the probe that was pre-equilibrated to 100 K. Sample temperature was inferred from the temperature of the stator and decreased from room temperature to 200 K in ~2 minutes followed by a slow decrease to 104 K over 15 minutes.

### Preparation of complex with nanodiscs

The nanodisc scaffold MSP1E3D1 was expressed and purified as previously described (Violeti R. et al 2020). POPG (1-palmitoyl-2-oleoyl-sn-glycero-3-phosphoglycerol) was used to assemble the nanodiscs. Buffer used for the nanodisc samples was 20 mM sodium phosphate, 50 mM NaCl, pH 7.4. Nanodiscs and α-synuclein were mixed at equimolar ratio, and $D_2O$ was added to obtain an 88:12 $D_2O$:$H_2O$ ratio. The sample was concentrated to the final concentration of 100 µM α-synuclein. Depleted deuterated glycerol ($d_8$-$^{12}$C-glycerol) and AMUPol were added as the last step for a final composition of 15:75:10 for $d_8$-$^{12}$C-glycerol:$D_2O$:$H_2O$ (v/v/v) respectively with 6.8 mM AMUPol.

### Preparation of fibrils

A-syn was eluted from size exclusion in 50 mM sodium phosphate pH 7.4, 0.5 mM EDTA (Tuttle MD, 2016) and fibrilized for 2 weeks at 37 °C shaking in a thermoblock at 1000 rpm. Fibrils were collected by ultracentrifugation (30 min, 100,000 x g). Monomeric non-fibrillized protein was removed by resuspending the fibril in buffer with same components (50 mM sodium phosphate pH 7.4, 0.5 mM EDTA) but with a $H_2O$:$D_2O$ ratio of 25:75. Fibrils were pelleted again by ultracentrifugation at 100,000 x g and the wash procedure was repeated one more time. The pellet was then weighted and mixed with $d_8$-$^{12}$C-glycerol containing AMUPol at 4.17 mM to result in final 60% glycerol, 10 % $H_2O$, 30 % $D_2O$, and 2.5 mM AMUPol. Fibrils were packed into a rotor, capped, and stored at -80 °C until measurements.

### NMR spectroscopy

All dynamic nuclear polarization magic angle spinning nuclear magnetic resonance (DNP MAS NMR) experiments were performed on a 600 MHz Bruker Ascend DNP NMR spectrometer/7.2 T Cryogen-free gyrotron magnet (Bruker), equipped with a $^{1}$H, $^{13}$C, $^{15}$N triple-resonance, 3.2 mm low temperature (LT) DNP MAS NMR Bruker probe (600 MHz). For $^{13}$C cross-polarization (CP) MAS experiments, the $^{13}$C radio frequency (RF) amplitude was fixed at 60 kHz and an $^{1}$H RF amplitude was 72 kHz. The 90° $^{1}$H pulse was 100 kHz, the 90° $^{13}$C pulse was 62.5 kHz, and $^{1}$H TPPM at 85 kHz for decoupling with phase alternation of ± 15° during acquisition of $^{13}$C signal. $^{13}$C-$^{13}$C 2D correlations were measured using 5 ms or 20 ms DARR mixing with the $^{1}$H amplitude at the MAS frequency. A total of 280 complex points in the indirect dimension were recorded with an increment of 25 µs. DARR experiments were apodized with a Lorenzto-Gauss window function with IEN-to-GB ratio of 2.5 and the IEN between 20 and 80 for both the $t_1$ and $t_2$ time domains.

### Construction of statistical coil ensemble and chemical shift predictions

The statistical coil ensemble of α-synuclein and the host-guest $Ala_{25}$-X-$Ala_{25}$ polypeptides, were constructed using flexible meccano algorithm (Ozenne et al. 2012, Salmon et al. 2009). In total, 2000 conformers were generated based on the α-synuclein sequence. Side chains were added using Rosetta Fixbb algorithm and structures were relaxed using Rosetta Relax (Alford R. et al. 2017). Chemical shift predictors PPM (Li D.-W., Brüschweiler R., 2012), SPARTA (Shen Y., Bax A., 2007), and SPARTA+ (Shen Y., Bax A., 2010) were used to predict chemical shifts for each amino acid in each conformer. One structure at a time was used as the input for the predictors (as opposed to an ensemble prediction) and the chemical shift predictions were performed using the default parameters. Phi-psi angles for each residue in the ensemble were extracted from each PDB of the ensemble using the Biopython package (Cock P. J. A., et al. 2009). Following this protocol, each residue in α-synuclein sequence has 2000 chemical shift predictions and 2000 phi/psi pairs. The predicted 1D spectrum representing a residue in the frozen ensemble was calculated by generating a Gaussian curve with a specified linewidth (default 1 ppm FWHM) for each of the 2000 chemical shift predictions. In case of multiple occurrences of an amino acid type (e.g., 10 threonines in the α-synuclein sequence), the final peak is a sum over all of them analogously to what we observe in the DNP MAS NMR spectra. The 2D peak shapes were generated from the chemical shift predictions using the histogram2d function of the NumPy python library and the contour function of the Matplotlib Python library (Harris CR, et al. 2020, Hunter JD 2007). Binning of 0.5 ppm was used for 2D contour plots of the 2D spectra. For the prediction of the 2D spectra of the threonine $C^{\beta}$-$C^{\gamma}$, each conformation was assigned a chemical shift for the $C^{\gamma}$ value that was randomly sampled from a normal distribution of $C^{\gamma}$ chemical shifts using the random.normal function from the NumPy library with the average (21.55 ppm) and standard deviation (0.5 ppm) of the values for threonine $C^{\gamma}$ described in the BMRB.

### Prediction of chemical shifts for fibril and membrane bound forms of α -syn

To predict the chemical shifts of the fibril core, we used the PDB entry 2N0A (Tuttle MD, 2016) of α-synuclein fibrils. Chemical shifts for the core were predicted from a single structure/chain while the disordered 'tails' were predicted separately relying on a flexible meccano fully disordered ensemble as described above. The predicted chemical shift values for atoms in the disordered "tails" and atoms of the fibril core were then combined, and each value was represented with a gaussian as described above. Before summing up the calculated gaussians to result in the final predicted spectrum, the simulated gaussians were scaled to adjust for the ensemble size (i.e., a single prediction for the atoms of the fibril core compared to 2000 predictions for an atom in an ensemble). It is worth noting that for proteins that contain both disordered and ordered regions, the contribution of the ordered residues dominates, due to the static disorder of the disordered regions and their consequently broad peaks.

To predict the chemical shifts for the "membrane bound" helical form of α-synuclein, we created an ideal helix geometry ($\phi$ = -60°, $\psi$ = -45°) for residues 1 to 93 using PeptideBuilder Python library (Tien MZ et al. 2013). The residues 1 to 93 were selected because they were previously described to form a helix (). Side chains were added using Rossetta fixbb and the structure was relaxed using Rosetta relax. The chemical shift predictions for the helical region were then combined with the predictions for the disordered 'tail' that was separately sampled using flexible meccano as described above.
