## Supplemental Figures 1-11 for "The conformational ensemble of an intrinsically disordered protein explains peak shapes under DNP conditions"

Supporting figures (11)

SI FIGURE S1

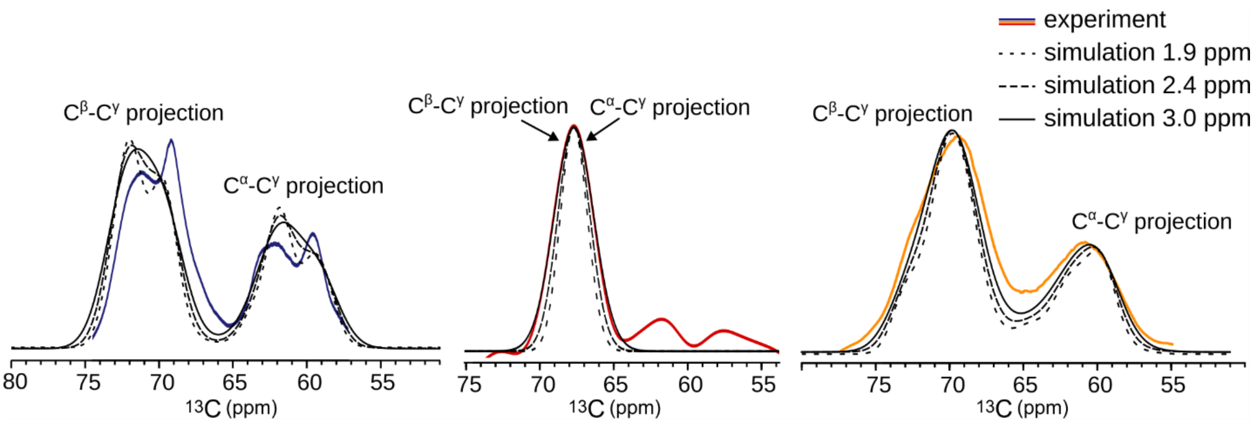


**Figure S1: Comparison of projections of experimental peaks with the simulated peak shapes with different broadening factor.** The experimental peak projections are indicated in colors: fibril (blue), nanodisc-bound (red), and disordered monomer (yellow).

SI FIGURE S2

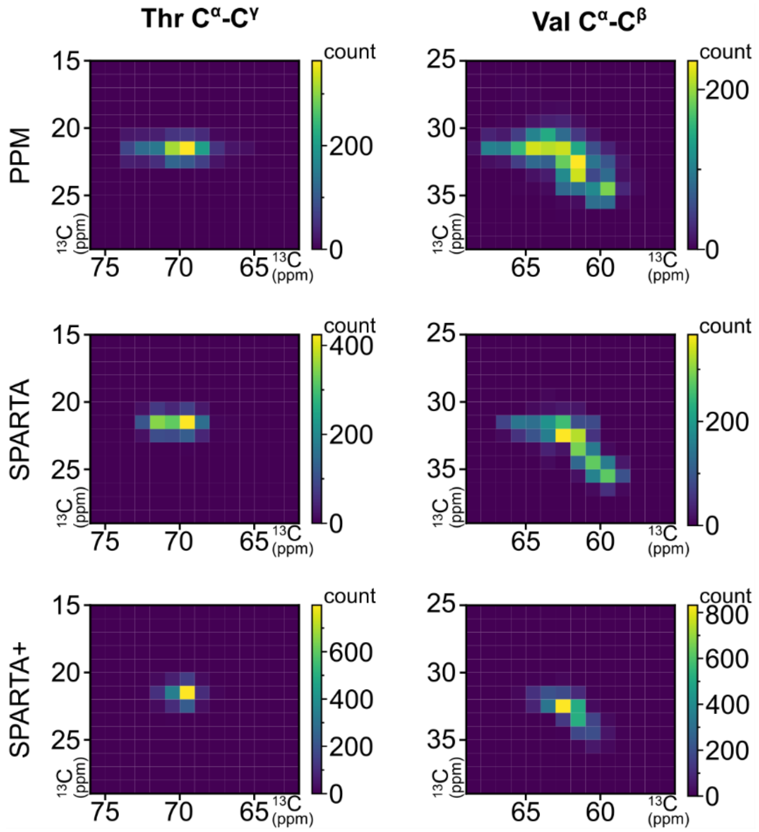


**Figure S2: Comparison of chemical shift predictions calculated using PPM_One, SPARTA, and SPARTA+.**

### SI FIGURE S3

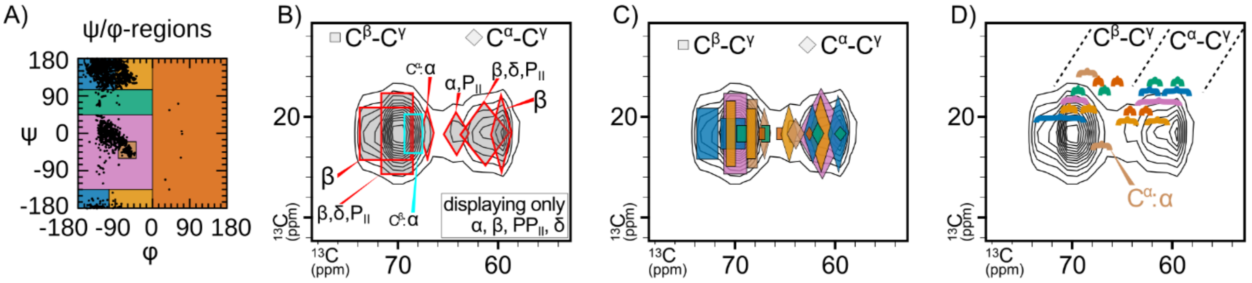


**Figure S3: Annotation of simulated threonine $C^{\beta}$-$C^{\gamma}$ and $C^{\alpha}$-$C^{\gamma}$ correlations.** A) Color definitions of the ψ/ϕ-regions. B) Simplified annotation that does not show the $\alpha_L$, ζ, and γ’ regions. C) All regions from (A) indicated on the simulated peak shapes. D) Alternative notation, with all the regions from (A) that highlights how the $C^{\alpha}$ chemical shift of conformers in α-helical conformation overlaps with the peak that is dominated by the $C^{\beta}$-$C^{\gamma}$ correlations.

SI FIGURE S4

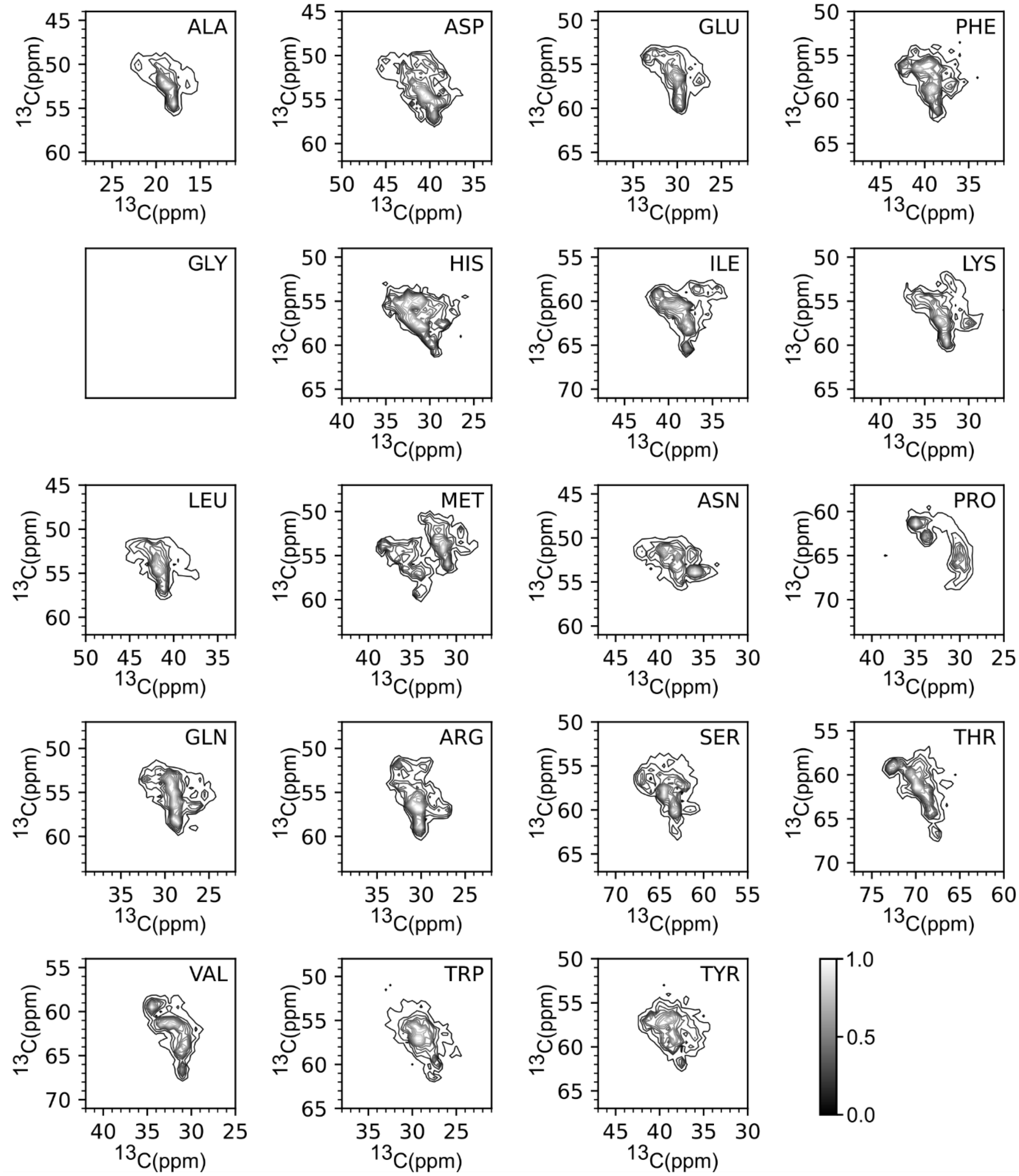


**Figure S4: Peak shape predictions for $C^{\alpha}$-$C^{\beta}$ correlations (bottom of the diagonal in a $^{13}C$-$^{13}C$ DARR-like spectrum).**

SI FIGURE S5

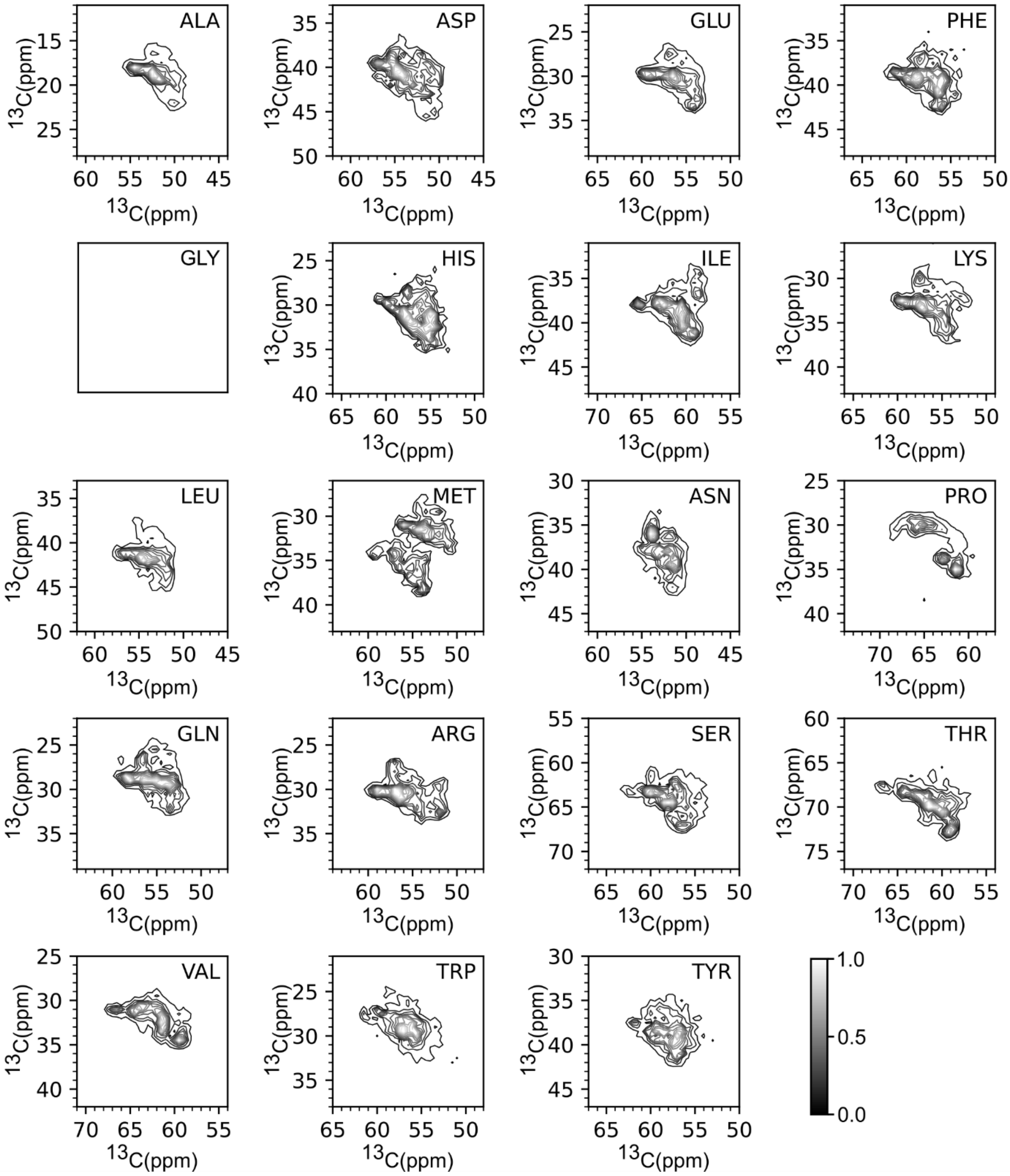


**Figure S5: Peak shape predictions for $C^{\alpha}$-$C^{\beta}$ correlations (top of the diagonal in a $^{13}C$-$^{13}C$ DARR-like spectrum).**

SI FIGURE S6

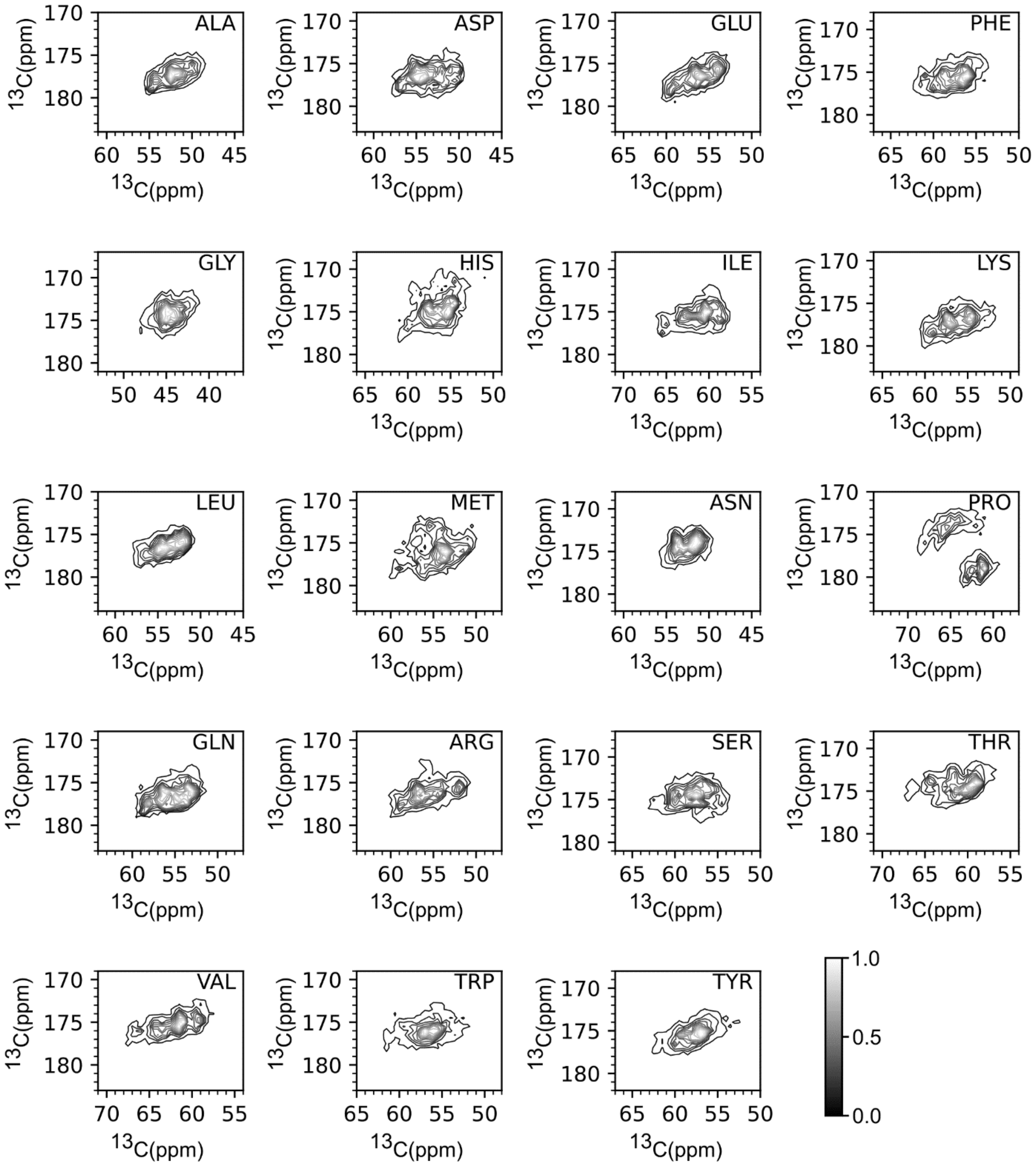


**Figure S6: Peak shape predictions for C'-C$^{\alpha}$ correlations (bottom of the diagonal in a $^{13}$C-$^{13}$C DARR-like spectrum).**

SI FIGURE S7

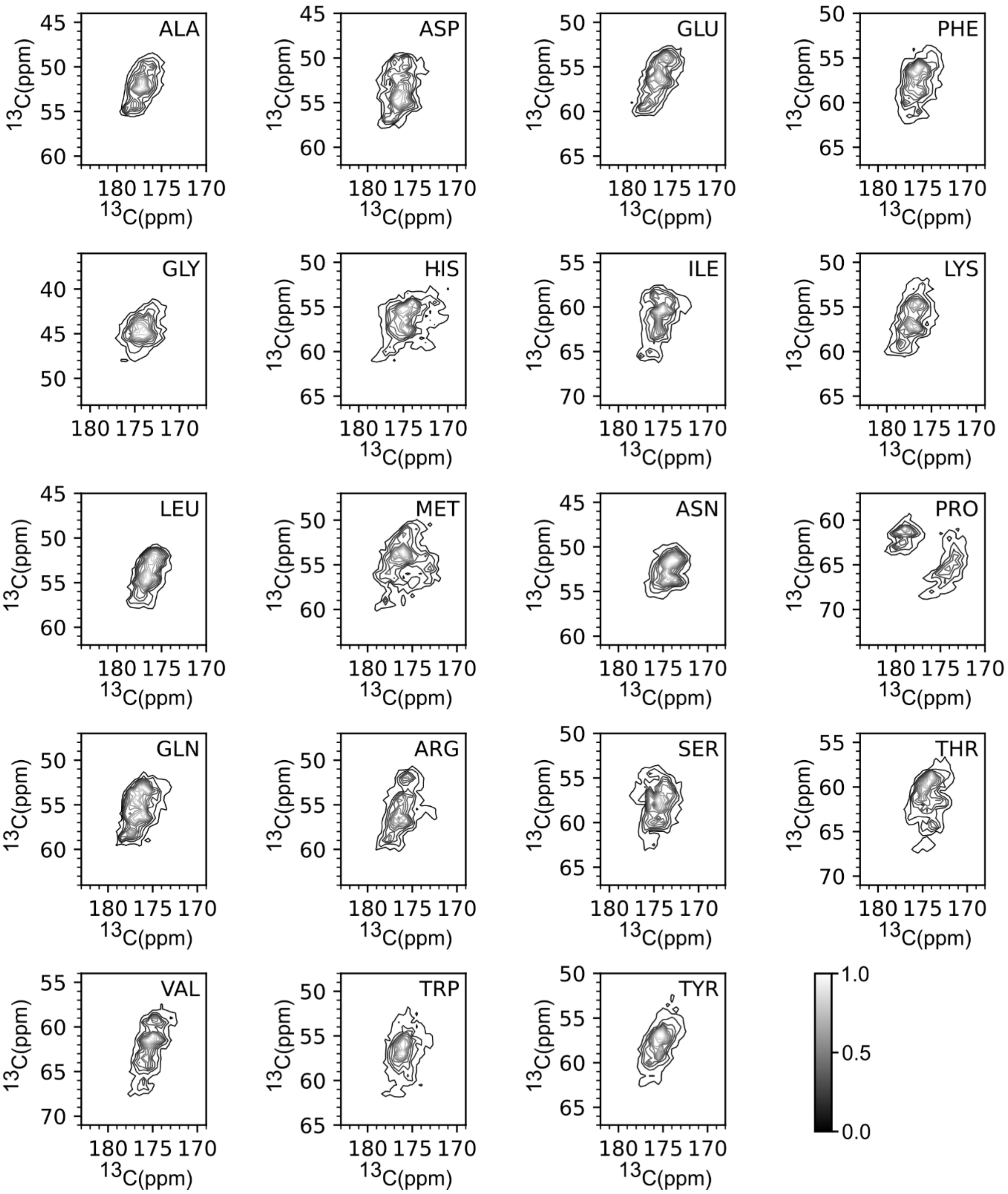


**Figure S7: Peak shape predictions for C'-C$^{\alpha}$ correlations. (top of the diagonal in a $^{13}$C-$^{13}$C DARR-like spectrum).**

SI FIGURE S8

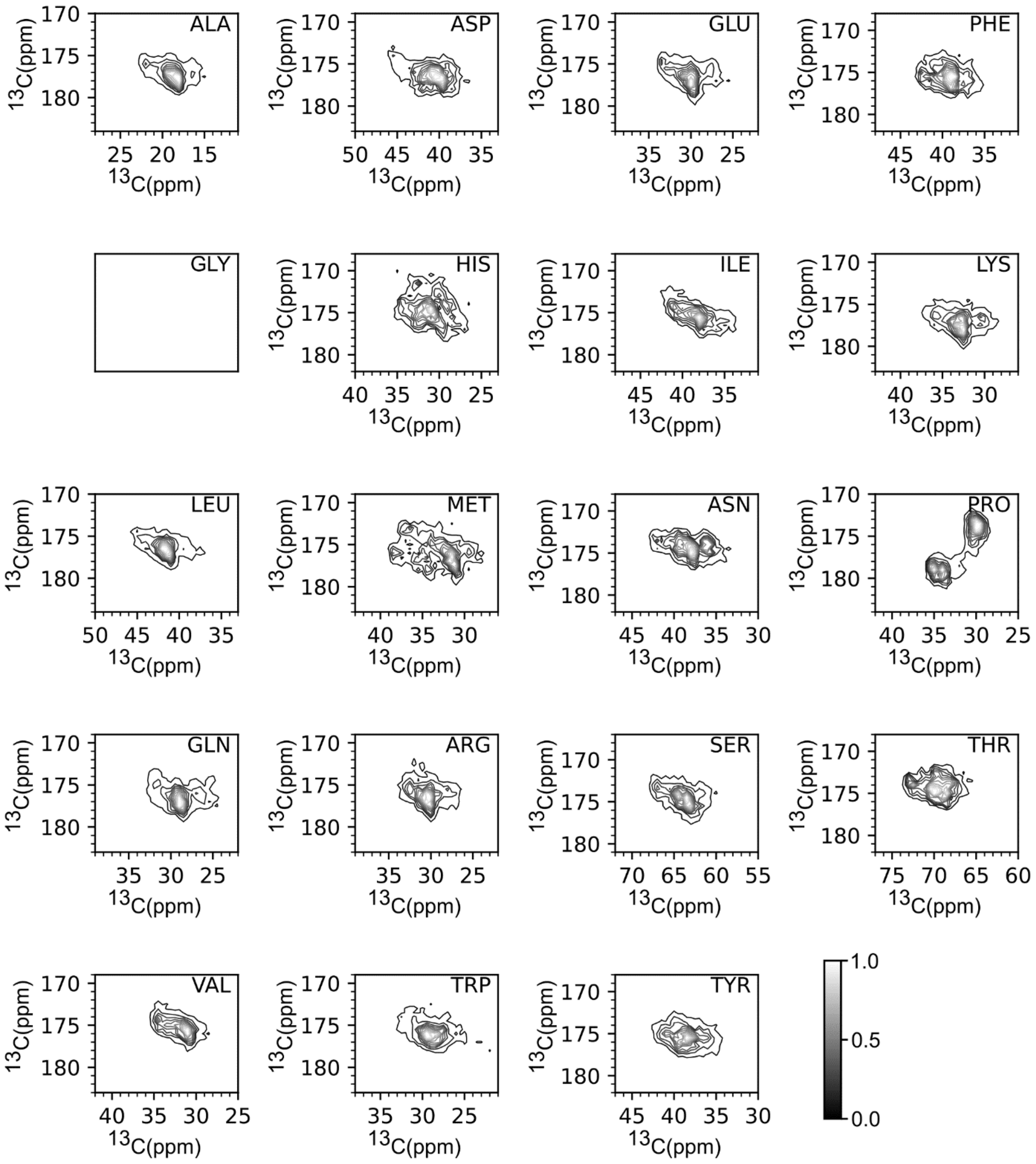


**Figure S8: Peak shape predictions for C'-Cβ correlations (bottom of the diagonal in a $^{13}$C-$^{13}$C DARR-like spectrum).**

SI FIGURE S9

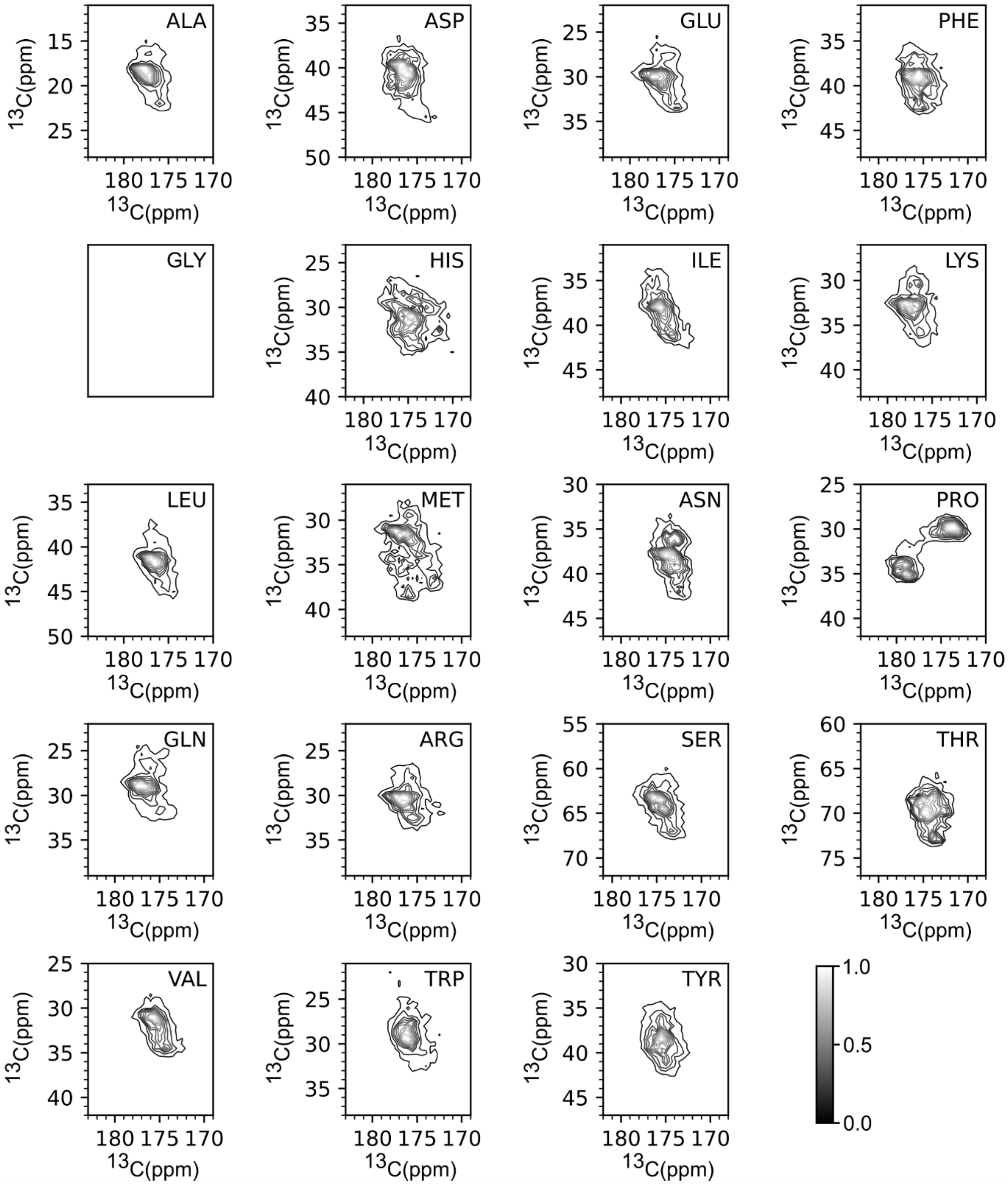


**Figure S9: Peak shape predictions for C'-Cβ correlations. (top of the diagonal in a $^{13}$C-$^{13}$C DARR-like spectrum).**

SI FIGURE S10

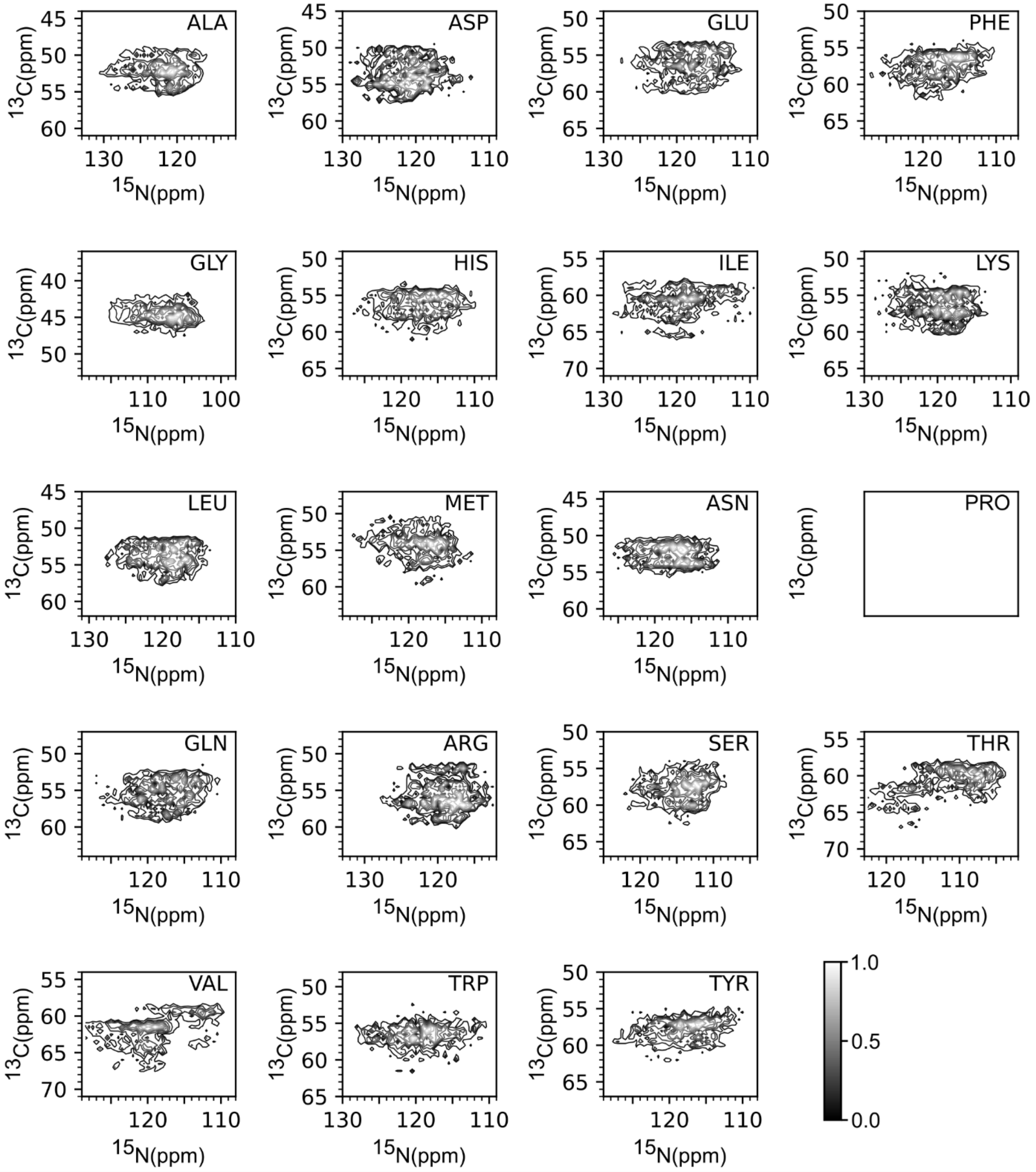


**Figure S10: Peak shape predictions for N-C$^{\alpha}$ correlations.**

SI FIGURE S11

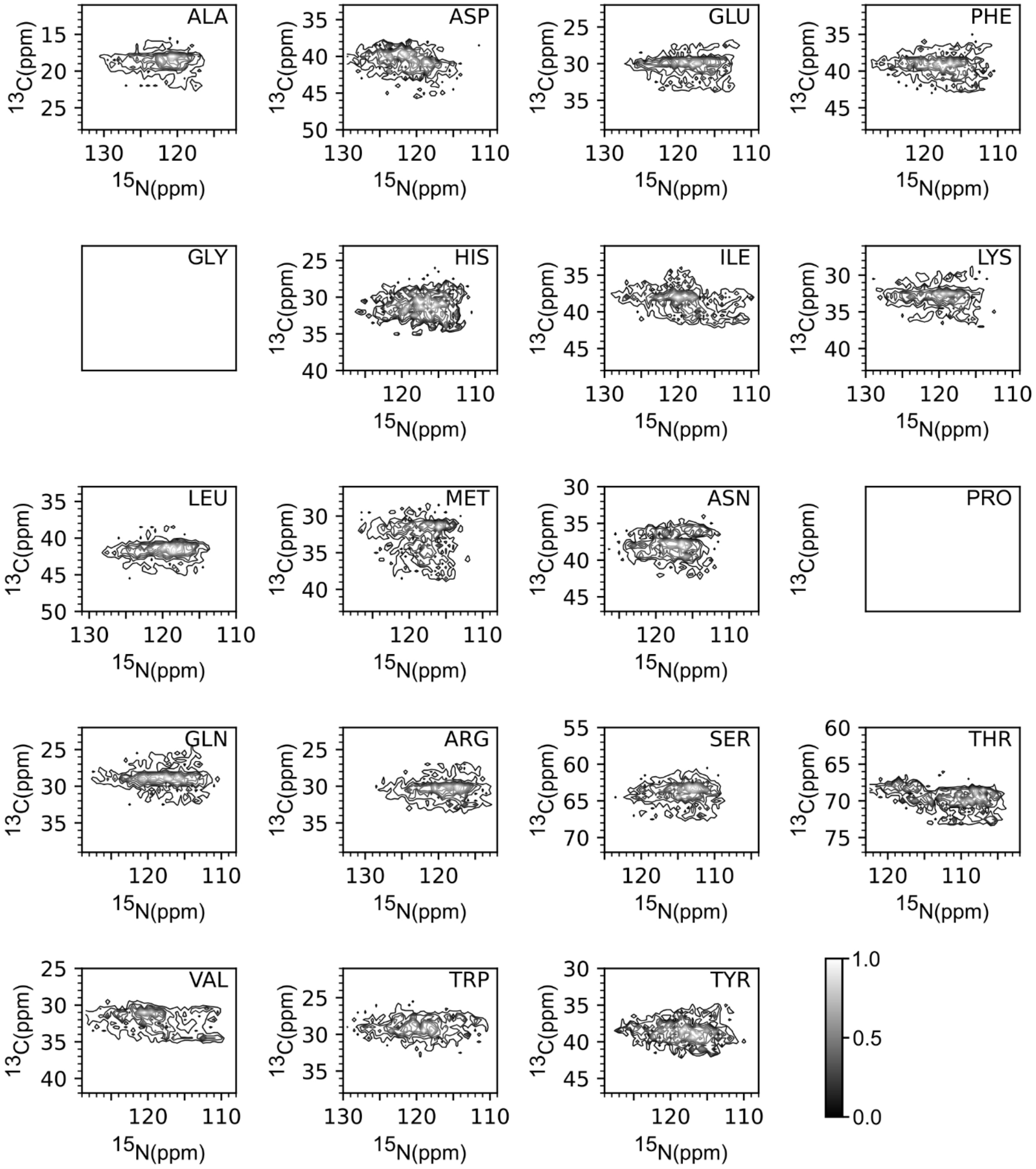


**Figure S11: Peak shape predictions for N-C$^{\beta}$ correlations.**
